# Transcriptomic analysis of non-model Drosophilidae reveals novel AMP candidates

**DOI:** 10.1101/2025.06.06.658223

**Authors:** Pankaj Dhakad, Dhobasheni Newman, Darren J. Obbard

## Abstract

**Background:** *Drosophila melanogaster* has been a valuable model for dissecting the molecular architecture of innate immunity. However, the family Drosophilidae encompasses over 4000 species, spanning deep evolutionary divergences and diverse ecologies. Here, we use immune challenge with the gram-negative pathogen *Providencia rettgeri* to investigate the conservation and evolution of immune responses in three non-model drosophilid species that diverged from *D. melanogaster* over 45 million years ago—*Hirtodrosophila cameraria*, *H. confusa*, and *Scaptodrosophila deflexa*.

**Results:** We find that all three species retain a core set of immune signaling and recognition genes, but exhibit substantial variation in effector gene content and inducibility. In particular, *Scaptodrosophila deflexa* lacks orthologs of multiple antimicrobial peptides (AMPs) known from *D. melanogaster*, including *DptA*, *AttA*, and *AttC*, and shows little transcriptional response to bacterial-challenge with *Providencia rettgeri*. In contrast, both of the *Hirtodrosophila* species exhibit substantial transcriptional responses, including strong induction of canonical Imd pathway genes. Microbiome profiling of our samples revealed higher *Providencia* abundance in *H. cameraria*, and high levels of the defensive symbiont *Spiroplasma* in *S. deflexa*—potentially explaining differences in infection outcome. Our combined annotation and expression analysis of these species also allowed us to identify 20 novel AMP-like candidates, many with structural features like known AMPs.

**Conclusions:** Our study demonstrates the feasibility of functional immune analyses in non-model *Drosophila* species and reveals striking lineage-specific differences in immune gene repertoire and expression. These findings highlight the importance of non-model, wild-derived taxa for uncovering novel immune effectors and understanding evolutionary forces shaping insect immunity.

## Background

In all organisms, the innate immune system forms the first line of defense against pathogens and parasites [1–3]. By rapidly recognizing and responding to infections, it reduces pathogen survival and replication—decreasing pathogen fitness to the benefit of the host. These antagonistic interactions often lead to coevolutionary dynamics, where immune-related genes—particularly those involved in pathogen recognition and effector functions—evolve at significantly higher rates compared to the genome-wide background [e.g. 4, 5-8]. The fruit fly *Drosophila melanogaster* has long served as a model for dissecting innate immunity in insects and helped in understanding key pathways such as Toll, Imd, JAK/STAT, and RNA interference (RNAi), which orchestrate defense responses against bacteria, fungi, and viruses [e.g. 9, 10, 11].

As in vertebrates, *D. melanogaster* mounts both humoral and cellular innate immune responses to combat pathogen infection. In *Drosophila*, cellular immunity primarily involves phagocytosis by plasmatocytes and encapsulation by lamellocytes, targeting invading microbes and larger parasites, respectively [12, 13]. In contrast, the humoral response mainly leads to the rapid production and systemic release of antimicrobial peptides (AMPs), which are secreted into the hemolymph to directly kill pathogens. Two major signaling pathways, Toll and Imd, regulate the majority of immune genes in *Drosophila*, including the production of AMPs. The Toll pathway is predominantly activated by lysine (Lys)-type peptidoglycan found in Gram-positive bacteria, as well as fungal β-glucans and host-derived microbial proteases. The Imd pathway is activated through the detection of diaminopimelic acid (DAP)-type peptidoglycan from Gram-negative and certain Gram-positive bacteria [11].

In addition to Toll and IMD, the JAK/STAT pathway plays a modulatory role in immune defense. While its primarily role is in development, stress response, and stem cell maintenance, it also contributes to immunity by regulating the expression of genes such as those encoding thioester-containing proteins (TEPs) and Turandot stress proteins [14–16].

The completion of 12 *Drosophila* genomes in 2007 opened the door to evolutionary analyses across multiple species, enabling researchers to investigate gene copy number variation, patterns of positive selection, and lineage-specific immune responses [17]. One striking pattern that emerged was an apparent dichotomy in the evolutionary dynamics of *Drosophila* immune genes; while core signaling components of immune pathways are often deeply conserved, the evolution of recognition and effector genes can be highly dynamic [8, 18, 19]. Upstream signaling molecules such as Relish and Dif family members typically persist as 1:1 orthologs, even in distantly related insects [20], and have detectable sequence homology in mammals (e.g., NF-κB), highlighting the deep conservation of immune signaling genes [21]. In contrast, AMP gene families frequently undergo duplication, pseudogenization, and loss, and exhibit high copy number variation between species [8, 19, 22, 23]. In some species, AMPs such as Drosocin, Drosomycin, Turandot, and Metchnikowin are either completely absent or show such a high sequence divergence that they are difficult to identify through standard homology searches [23–25].

This contrast raises an important but hard to answer question: what drives the dynamic evolutionary patterns of antimicrobial peptides (AMPs)—including gene duplications, losses, and lineage-specific expression? This question is difficult to answer, in part, because our experimental understanding of the drosophilid antibacterial response comes largely from a few of the more experimentally tractable species, predominantly *D. melanogaster* and its close relatives within the subgenus *Sophophora* [26]—with only handful of species outside of this subgenus [18, 23–25, 27]. Nevertheless, comparative studies have revealed striking lineage-specific variation in effector gene repertoires.

The antimicrobial peptide *diptericin* offers a particularly well-characterized example of this variation. *Diptericin* genes are key effectors of the Imd pathway and show differential inducibility by Gram-negative bacteria across *Drosophila* species [23]. A population genetics study found that serine/arginine polymorphism at 69^th^ residue of *DptA* significantly alters susceptibility to *Providencia rettgeri* bacterial infection in *D. melanogaster* and *D. simulans*– an association interpreted as a signature of balancing selection [28]. More recent work has shown that the selective landscape acting on *diptericin* may be highly context-dependent: interactions between host genotype, sex, environmental stress (e.g., starvation), and pathogen exposure can shape the evolutionary trajectories of AMP alleles, potentially maintaining diversity over time [29].

At a broader phylogenetic scale, AMP families exhibit even more extreme evolutionary dynamics. For example, while *DptA* has been lost or pseudogenized in some non-melanogaster species, others possess divergent paralogs (e.g., *DptC* in the subgenus *Drosophila*) that are syntenic but highly diverged at the sequence level [18]. Consistent with this, AMP duplicates can undergo neofunctionalization: in *D. virilis*, a *defensin* paralog has evolved from a role in bacterial killing to one in toxin neutralization [30]. Together, these findings suggest that while the basic architecture of innate immunity is ancient and broadly conserved, the downstream effector repertoire is evolutionarily labile and shaped by selection from species-specific microbial exposure, life-history, and ecological pressure.

Now, with the recent availability of nearly 400 drosophilid genomes [31]—over 300 of which are annotated [32]—there is an opportunity to explore the extent to which canonical immune responses are conserved across the drosophilid phylogeny. For example, we can ask if deeply diverged lineages harbor novel immune effectors, or if more distantly diverged non-model species—with their unique ecological niches and evolutionary histories—possess a distinct immune repertoire. However, one major challenge remains; most functional studies have focused on species amenable to long-term laboratory culture, but these represent only a small fraction of drosophilid diversity [31]. Understanding gene function in species that are not easily cultured in large numbers remains challenging.

In this study, we demonstrate the feasibility of characterizing immune responses to bacterial-challenge in non-model, less easily cultured, species of Drosophilidae by performing comparative transcriptomic analysis on individual first-generation wild-derived flies. We selected three common European species for analysis—*Hirtodrosophila cameraria*, *Hirtodrosophila confusa*, and *Scaptodrosophila deflexa—*each of which is highly divergent from *D. melanogaster* and other well-studied taxa [>45 Mya; Fig. 1; 33]. These species are not only genetically distant, but also ecologically distinct. *Hirtodrosophila confusa* is a relatively large drosophilid and a fungal specialist that thrives in cool temperate environments and is frequently found in association with large fungal fruiting bodies such as dryad’s saddle (*Polyporus squamosus*). *Hirtodosophila cameraria* is also a specialist fungus breeder, moderately abundant in the UK on basidiomycete fungi such as *Phallus impudicus* and *Lactarius quietus* [34, 35]. *Scaptodrosophila deflexa*, in contrast, is thought to lay its eggs in yeast-rich sap fluxes. However, despite their broad distribution and ecological interest, these species remain very poorly studied; for example, the genera *Hirtodrosophila* and *Scaptodrosophila* have both been shown to be polyphyletic [Hirtodrosophila partly within the paraphyletic Drosophila; 31, 36]. Despite the availability of high-quality genome sequences [31, 34], they are yet to receive any attention in functional or comparative genomics. To our knowledge, only a single report addresses egg viability in *H. confusa* [37], and no studies to date have explored their immunogenomics or transcriptomic responses to infection. By analyzing these lineages, we aim to expand our understanding of immune system diversity and evolution within Drosophilidae, moving beyond the traditional model organisms and exploring a broader phylogenetic landscape.

**Fig. 1.**
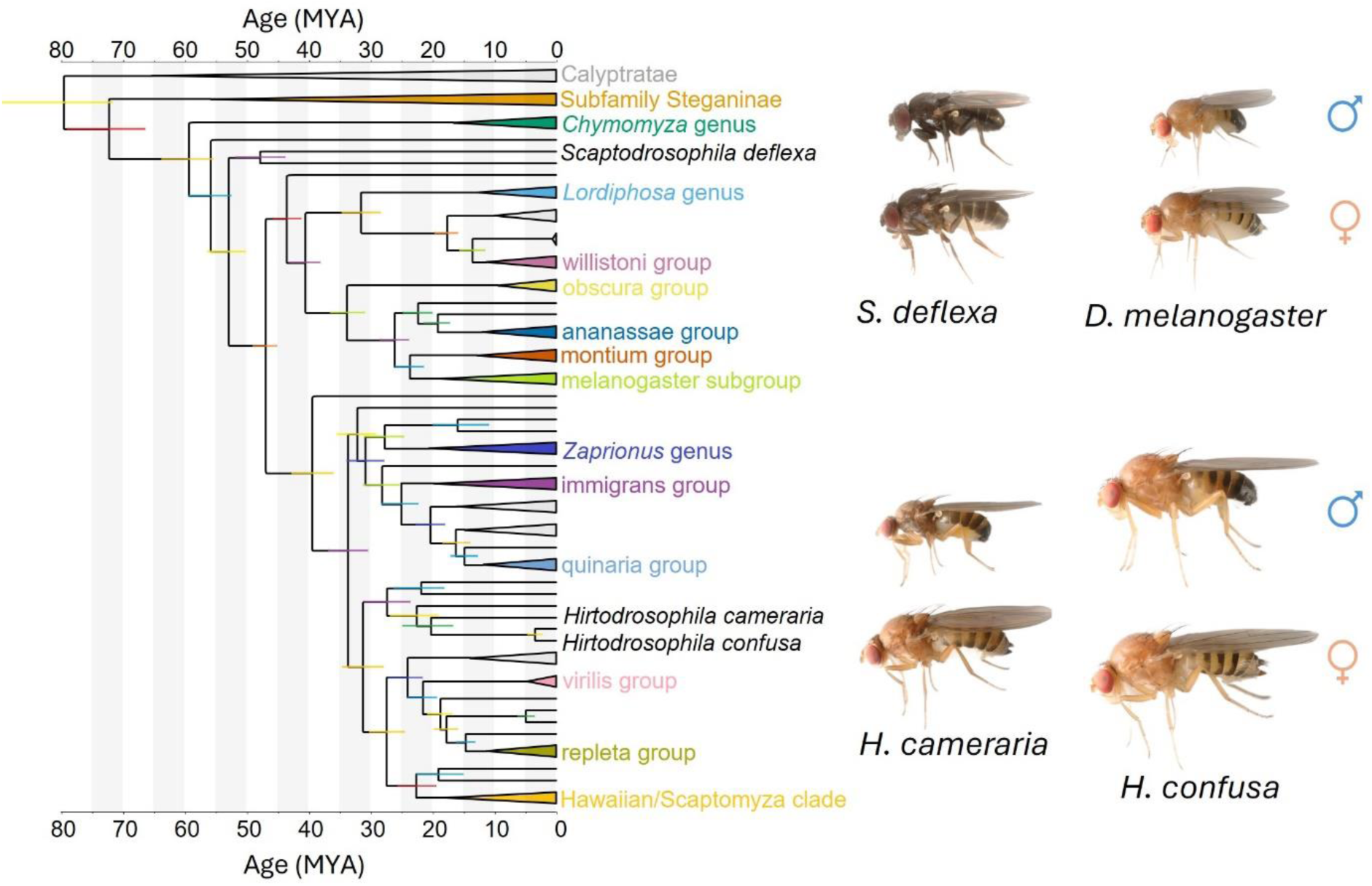
The phylogenetic position of the study species within Drosophilidae. An approximate time-calibrated phylogeny showing the relationships of major lineages of Drosophilidae, including the placement of the three non-model species used in this study— *Hirtodrosophila cameraria*, *H. confusa*, and *Scaptodrosophila deflexa*. The *Hirtodrosophila* species belong to a diverse and understudied cluster within the subgenus *Drosophila*, while *S. deflexa* branches deep within the subfamily Drosophilinae, representing one of the earliest branching lineages relative to *D. melanogaster* (mrca >50 MYA). Divergence times are shown in millions of years ago (MYA) along the x-axis. Taxonomic groups are color-coded by genus/subgenus or species group, and collapsed clades indicate major drosophilid lineages for clarity. Images to the right right show adult male (♂) and female (♀) flies of each of the three focal species, plus *D. melanogaster* at the same scale for comparison. The tree figure was generated as described in Dhakad P et al. [32], based on 285 single-copy BUSCO orthologs.

Here we compare the transcriptomes of unchallenged flies with those of flies we experimentally challenged with the Gram-negative bacterial pathogen *Providencia rettigeri*. Our goals are to: (i) evaluate whether pathogen-challenged transcriptomes improve the annotation of immune genes in species lacking reference-quality genomes; (ii) assess the feasibility of differential expression analyses in these non-model species; and (iii) identify conserved and novel immune-related genes, including potential AMPs. Our findings suggest that despite their deep evolutionary divergence, bacterial infection induces measurable and broadly similar transcriptional responses in *H. cameraria* and *H. confusa* species but not in the more distant *S. deflexa*—a pattern that could potentially be shaped by differences in microbiota, immune strategy, or symbiont-mediated protection.

## Results

### Pathogen challenge does not substantially improve annotation

To assess the impact of the availability of pathogen-challenged RNAseq on gene predictions, we generated three separate genome annotations for each species using BRAKER3 [38]. The three annotations were based on (i) RNA-seq from pathogen-challenged flies, (ii) RNA-seq from unchallenged flies, and (iii) combined data. All three annotations recovered the same set of core genes in each species, and most gene models were shared between the challenged and unchallenged annotations, with only a small number unique to one set or the other (Table 1; Fig. 2). The combined RNA-seq data produced slightly more gene models compared to either pathogen-challenged or unchallenged datasets alone for *H. cameraria* and *S. deflexa*, whereas the pathogen-challenged dataset yielded slightly more genes for *H. confusa* (Table 1). To assess the global similarity between genome annotations generated from pathogen-challenged and unchallenged RNA-seq datasets, we counted the shared (i.e. overlapping coordinates) and unique gene models, recording how many were assignable to an orthogroup (Fig. 2). As expected, most genes were shared between pathogen-challenged and unchallenged annotations (category C; Fig. 2), suggesting consistent annotation of core gene sets across datasets. Only a small number of genes were uniquely recovered in either pathogen-challenged or unchallenged annotations (categories A, B, D, and E; Fig. 2), and many of these lacked orthogroup assignment, implying that they may be annotation errors, or potentially novel genes. To examine whether pathogen-challenged RNA-seq improved recovery of immune-related genes, we compared immune gene orthogroup [‘Hierarchical OrthoGroup’; HOG; 39] representation between pathogen-challenged and unchallenged annotations for each species. Small differences were observed, but most immune genes were either recovered in both annotations or missing from both, with few genes uniquely recovered by pathogen-challenged RNA-seq (Table 1). To complement this analysis, we performed Gene Ontology [GO; 40] enrichment analysis on genes uniquely annotated from pathogen-challenged or unchallenged RNA-seq datasets. However, no significant enrichment for immune-related terms was detected among genes uniquely recovered in pathogen-challenged annotations (Additional file 1). Overall, these results indicate that pathogen-challenged RNA-seq did not substantially improve overall immune gene discovery or annotation completeness compared to unchallenged RNA-seq.

**Table 1:** Gene annotation statistics and immune gene recovery from pathogen-challenged, unchallenged, and combined RNA-seq data for each species.

| Species | Genes | Mean CDS length (Kbp) | Immune OGs | Immune genes | Immune CDS length (Kbp) |
| --- | --- | --- | --- | --- | --- |
| *D. melanogaster* | 13986 | 1.54 | 568 | 638 | 1.74 |
| *H. cameraria* (unchallenged) | 14407 | 1.46 | 515 | 545 | 1.88 |
| *H. cameraria* (challenged) | 14203 | 1.47 | 520 | 551 | 1.86 |
| *H. cameraria* (combined) | 14503 | 1.47 | 539 | 573 | 1.87 |
| *H. confusa* (unchallenged) | 14200 | 1.39 | 490 | 519 | 1.72 |
| *H. confusa* (challenged) | 14469 | 1.39 | 500 | 530 | 1.78 |
| *H. Confusa*<br>(combined) | 14300 | 1.41 | 534 | 566 | 1.76 |
| *S. deflexa*<br>(unchallenged) | 13017 | 1.50 | 490 | 519 | 1.95 |
| *S. deflexa*<br>(challenged) | 12991 | 1.48 | 485 | 514 | 1.93 |
| *S. Deflexa*<br>(combined) | 13148 | 1.50 | 522 | 553 | 1.95 |

**Fig. 2:**
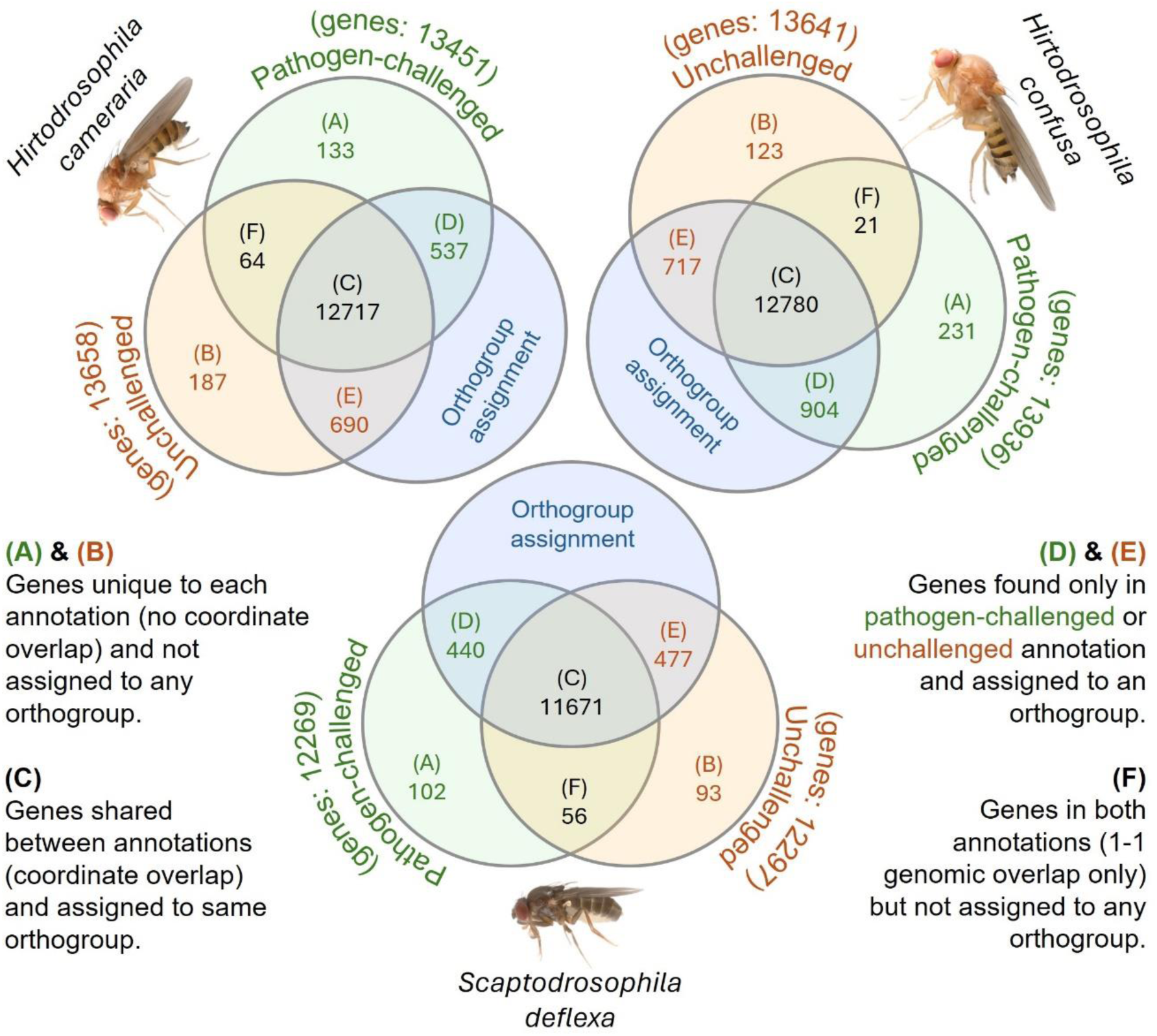
Comparison of gene sets annotated using RNA-seq from pathogen-challenged and unchallenged samples. Venn diagrams illustrating the overlap of gene models annotated using RNA-seq from pathogen-challenged (green) and unchallenged (orange) samples of *Hirtodrosophila cameraria*, *H. confusa*, and *Scaptodrosophila deflexa*. Gene sets were compared based on (i) genomic coordinate (1:1 overlap), and (ii) orthogroup assignment using OrthoFinder. Numbers inside Venn diagrams (A to F) represent gene model counts. The total number of predicted genes per condition is shown in parentheses next to each label.

### The detectable immune repertoire differs between species

To further assess how immune gene recovery varied across the three divergent drosophilid lineages, we examined the presence/absence of genes in *H. cameraria*, *H. confusa*, and *S. deflexa* that have homology with a curated set of 638 well-characterized *D. melanogaster* immune-related genes [26]. These 638 immune genes were clustered into 568 HOGs. Of these, *H. cameraria* recovered genes in 539 HOGs, *H. confusa* in 534, and *S. deflexa* in 522 (Table 1; Additional file 2). In total, 497 HOGs had immune genes recovered across all three species, while 11 HOGs contained only *D. melanogaster* genes. These uniquely missing HOGs included several AMPs such as *drosocin* (*Dro*), all members of the *turandot* family, and *drosomycins* (*Drs* and *Drsl*1–6; Additional file 2). These apparent losses support previous observations that the *drosomycin* and *turandot* AMP families are largely restricted to *D. melanogaster* and closely related species in the subgenus *Sophophora* [18, 25]. However, we found homologs of a *drosocin*-like gene in all three species, and in *H. confusa* and *H. cameraria* this gene encodes multiple tandem repeats of the peptide domain—as previously reported in *D. neotestacea* and other species in the subgenus *Drosophila*. In total, only two recognition proteins (*PGRP-SB2* and *CG12780*) and six signaling genes (*serpin 42Dd*, *MstProx/Toll-3*, *Toll-4*, *sphinx1*, *sphinx2*, and *amnesiac*) were missing from all three species. Interestingly, *S. deflexa* lacked orthologs of *diptericin A* (*DptA*), *attacin C* (*AttC*) and *attacin D* (*AttD*), suggesting lineage-specific loss of these AMPs.

To test whether the species and immune-gene functional categories differed in the probability of each ‘gene’ (i.e. orthogroup) being recovered, we fitted a Bayesian binomial linear mixed-effects model using MCMCglmm. Posterior predictions revealed a significant species effect, with *H. cameraria* (posterior mean = 99.88%; 95% Credible Interval: 99.61-99.98) and *H. confusa* (99.79%; CI: 99.35-99.97) showing similar probabilities of immune gene recovery, while *S. deflexa* recovered significantly fewer immune genes (99.58%; CI: 98.80-99.93). This may partly reflect differences in genome and annotation quality, but also likely reflects genuine differences in immune gene conservation and loss. We also detected a marginal effect of gene category, such that canonical signaling genes (99.97%; CI: 99.91-99.99) were more likely to be recovered than effector genes (99.46%; CI: 98.23-99.94), although receptor and unknown categories were not significantly different to effectors. Among interaction terms, only the combination of *S. deflexa* and signaling genes showed a significant positive deviation from expectation (99.97%, CI: 99.89–99.99), indicating that signaling genes were relatively well retained even in *S. deflexa*. Other species and category interactions terms were not significantly different from additive expectations.

To illustrate the evolutionary lability of effector gene families, we chose to examine the *diptericin* and *attacin* genes in more detail. Phylogenetic analysis of *diptericins* (Fig. 3 A) confirmed three distinct clades: *DptA*, *DptB*, and *DptC*, the latter being restricted to the subgenus *Drosophila* [18]. *Scaptodrosophila deflexa* encoded only a single *DptB*-like gene, clustering with orthologs from *H. confusa*, *H. cameraria*, and *D. melanogaster*, but lacked any detectable *DptA* and *DptC* homologs. In contrast, multiple *DptA* and *DptC* paralogs were recovered in both *Hirtodrosophila* species, suggesting lineage-specific expansions. Similarly, the *attacin* gene tree (Additional file 3) showed that *H. cameraria* and *H. confusa* have multiple copies of *AttC*, and *AttA/B*, while *S. deflexa* possesses only a single *AttA/B*-like gene with no detectable orthologs of *AttC* and *AttD* (Fig. 3 C; Additional file 3). The clear pair of *AttA* and *AttB* genes in most species (Additional file 3) might superficially suggest that each species has experienced a recent duplication independently. However, it is more likely that these patterns reflect concerted evolution in *diptericin* and *attacin* gene families [19, 41], highlighting the challenges of inferring orthologs of AMPs across divergent genomes. Taken together, these results show the rapid and idiosyncratic evolution of effector gene repertoires across drosophilid species, and illustrate how their annotation is particularly sensitive to both sequence divergence and genome assembly quality.

**Fig. 3:**
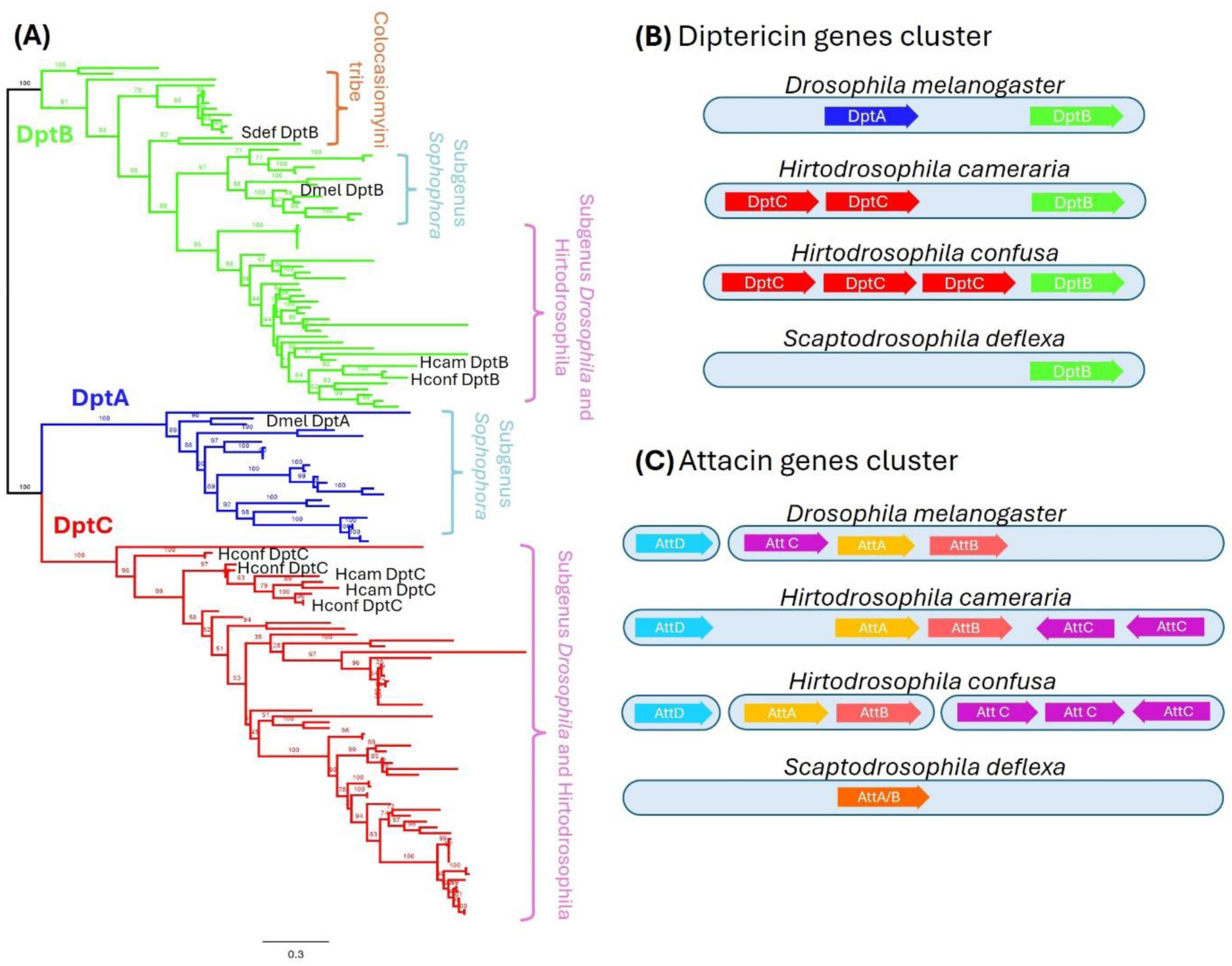
Divergence and lineage-specific organization of Diptericin and Attacin gene families in drosophilids. **(A)** Maximum-likelihood gene tree of *diptericin* genes from 51 drosophilid species, including *Hirtodrosophila cameraria*, *H. confusa*, and *Scaptodrosophila deflexa*, generated using IQ-TREE2 based on aligned amino acid sequences. The gene tree highlights three distinct clades corresponding to *DptA* (blue), *DptB* (green), and *DptC* (red). *DptC* is restricted to the subgenus *Drosophila*, including the *Hirtodrosophila* species, and is absent from both *D. melanogaster* and *S. deflexa*. **(B)** Synteny of *diptericin* gene clusters in *D. melanogaster*, *H. cameraria*, *H. confusa*, and *S. deflexa*. In *D. melanogaster*, DptA is upstream of DptB. In contrast, the *Hirtodrosophila* species contain multiple tandem repeats of *DptC* genes upstream of *DptB* inplace of *DptA*. *Scatopdrosophila deflexa* has only a single *DptB*-like gene and lacks both *DptA* and *DptC*, possible indicating lineage-specific gene loss or pseudogenization. **(C)** Synteny of *attacin* gene clusters in the same four species. In *D. melanogaster*, *attacin* genes are on two different chromosomes, AttD on 3R and AttC, AttA, and AttB on 2R. *Hirtodrosophila cameraria* exhibits all four *attacin* genes on same contig, with two *AttC* duplicates positioned downstream of *AttA* and *AttB*. *Hirtodrosophila confusa* also retains *AttD* and *AttC* (3 copies), although on different contigs. In contrast, *S. deflexa* has only a single *AttA/B*–like gene, with no identifiable orthologs of *AttC* or *AttD.* **Note:** Gene cluster diagrams in panels B and C are schematic and not drawn to scale; gene sizes and intergenic distances are illustrative only.

### Immune challenge triggers a conserved immune response in *Hirtodrosophila* but not in *S. deflexa*

To identify genes that are transcribed in response to bacterial infection, we performed differential expression analysis between pathogen-challenged and unchallenged individuals using DESeq2 [42]. We identified 363 significantly upregulated genes (adj. p < 0.05, log₂ fold change ≥ 1) in *H. cameraria*, 149 genes in *H. confusa* and only 34 genes in *S. deflexa* after bacterial challenge (Additional file 4). In addition, we identified 230 significantly downregulated genes (adj. p < 0.05, log₂ fold change ≤ -1) in *H. cameraria*, 82 genes in *H. confusa* and only 7 genes in S*. deflexa* (Additional file 4).

To visually assess the consistency of the response across individuals, we performed principal component analysis (PCA) of normalized expression data. In both *H. cameraria* and *H. confusa*, PCA analysis revealed clear separation between pathogen-challenged and unchallenged samples along PC1 (explaining 63% and 56% variance, respectively), confirming a strong and coordinated response to infection (Fig. 4 A). In contrast, samples from *S. deflexa* showed no clear separation by treatment, indicating limited transcriptional response and/or greater inter-individual variability (Fig. 4 A). Heatmaps of significantly differentially expressed genes corroborated these patterns, further highlighting the strong transcriptional induction in the two *Hirtodrosophila* species and the lack of a detectable response in *S. deflexa* (Fig. 4 B).

**Fig. 4:**
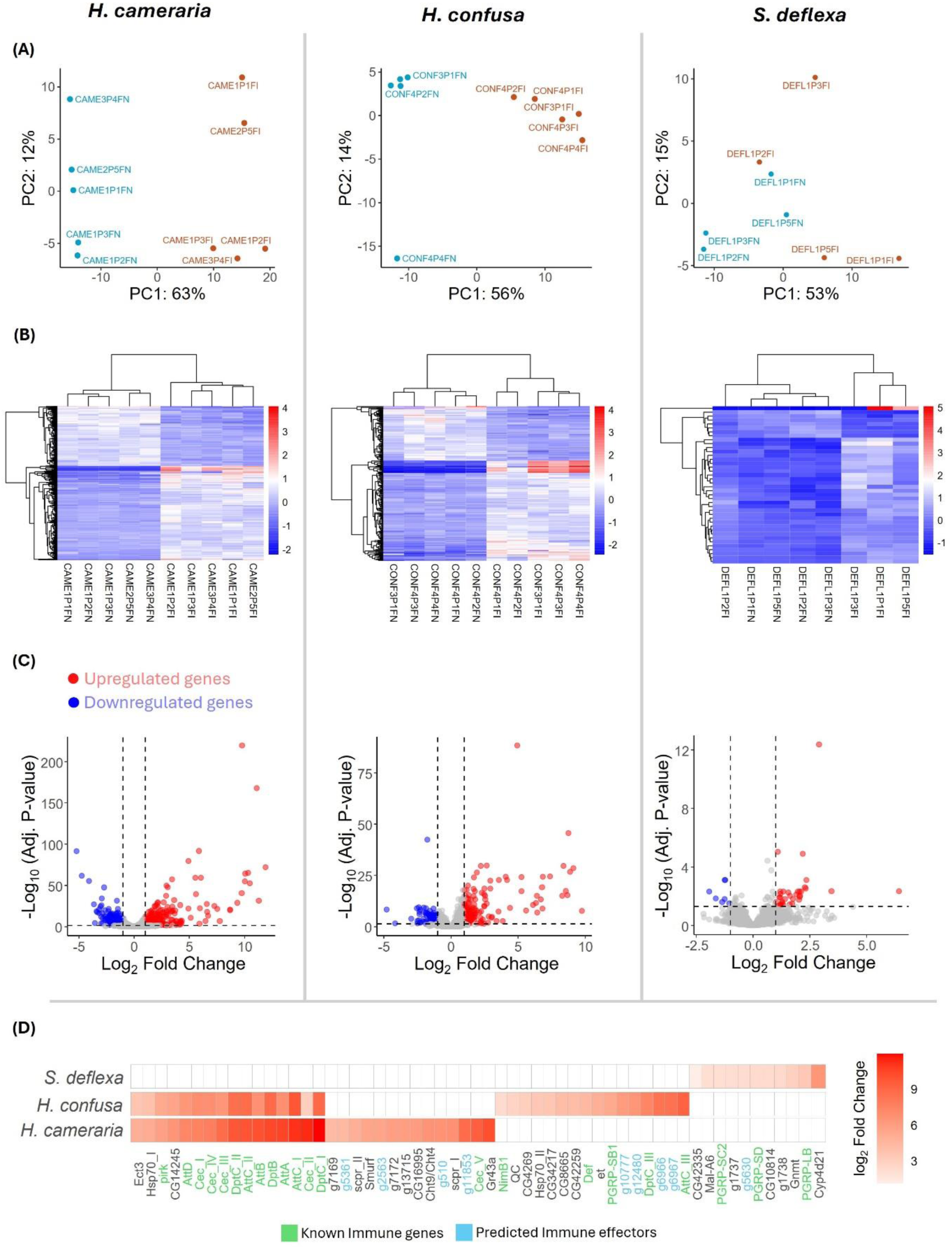
Pathogen-challenge with *Providencia rettgeri* induces immune responses in *Hirtodrosophila* species but not in *S. deflexa*. **(A)** Principal component analysis (PCA) of normalized gene expression data shows clear separation between pathogen-challenged and unchallenged individuals in *H. cameraria* and *H. confusa*, but not in *S. deflexa*, indicating weaker transcriptional response in the latter. **(B)** Heatmaps of significantly differentially expressed genes (adj. p < 0.05, |log₂FC| ≥ 1). **(C)** Volcano plots displaying log₂ fold change versus -log_10_ adjusted p-values for all expressed genes, highlighting significantly upregulated genes (red) and significantly downregulated genes (blue). **(D)** Heatmap showing log₂ fold change of selected top genes (based on log_2_ fold change and adj. p value) following pathogen-challenge across all three species. Canonical Imd-pathway target AMPs such as *diptericin*, *attacin*, and *cecropin* are strongly induced in *H. cameraria* and *H. confusa*, but not in *S. deflexa*. Known immune effectors are highlighted in green and novel immune effectors predicted in this study are highlighted in blue.

As expected from Gram-negative bacterial challenge, the most strongly upregulated genes in *H. cameraria* and *H. confusa* were homologs of canonical AMPs—such as *diptericins*, *attacins*, and *cecropins*—that are downstream targets of the Imd pathway in *Drosophila melanogaster* (Fig. 4 C and D). Interestingly, *bomanins*—targets of Toll pathway—were also upregulated, although upregulation was not to the level of Gram-negative specific AMPs. We also observed significant upregulation of peptidoglycan receptors, such as homologs of *PGRP-SB1*, *PGRP-LB*, *PGRP-SD*, *PGRP-SA*, *PGRP-LF*, *PGRP-LC*, and *PGRP-SC2* and as well as serine proteases (e.g. *sp7*), which have established roles in immune activation and microbial killing (Fig. 4 D; Additional file 4). Apart from immune effectors, signaling genes such as *Rel* and *pirk* that regulate the Imd pathway were also upregulated. In contrast, *S. deflexa* only showed detectable induction of three PGRP receptors among known immune genes (*PGRP-LB*, *PGRP-SD*, and *PGRP-SC2*), consistent with a weak or absent transcriptional immune response to *Providencia rettgeri* (Fig 4 D; Additional file 4).

Despite the overall similarity of the transcriptional response to infection in *H. confusa* and *H. cameraria*, notable differences may nevertheless exist. For example, *defensin* (*Def*) and *IM33* were only detectably induced in *H. confusa,* and *transferrin 1* and *listericin* only detectably induced in *H. cameraria* (Fig 4 D; Additional file 4). Additionally, in all species, we identified several strongly induced genes without clear homologs in *D. melanogaster*, suggesting the induction of novel or species-specific immune genes, many of which might be novel AMPs (below).

We carried out Gene Ontology analyses of the differentially expressed genes to help identify functional classes of genes whose expression is induced or repressed upon pathogen challenge. In both *H. cameraria* and *H. confusa*, upregulated genes were significantly enriched for terms such as “defense response to Gram-negative/Gram-positive bacteria”, “antibacterial humoral response” and “response to fungus” (Additional file 5). These enrichments confirm that infection induced a coordinated immune response. In contrast, only a few immune-related GO terms (“defense response to Gram-negative bacteria” and “antibacterial humoral response”) were significantly enriched among the few upregulated genes in *S. deflexa*, consistent with its subdued transcriptional response. Downregulated genes across all species were enriched for terms related to metabolism and structural processes.

### Microbiome variation could underlie immune response heterogeneity

All flies were first-generation lab-reared individuals derived from wild-caught parents, and were reared on non-standard media. This increases the likelihood that the flies carry diverse (and divergent) microbial communities. Such microbiota can influence baseline immune status, alter pathogen susceptibility, or modulate the host’s immune response to bacterial-challenge [43–46]. Thus, to explore whether microbiome composition varies, and might therefore have contributed to the apparent variation in immune responses, particularly the muted transcriptional induction observed in *S. deflexa*, we profiled the taxonomic abundance using the unmapped RNA-seq reads.

We first constructed a *de novo* assembly of unmapped reads for each library, followed by ‘diamond blastp’ [47] searches against the NCBI ‘nr’ protein database for taxonomic assignment. The assembly of unmapped reads produced between 15,439 and 45,168 contigs per sample, and taxonomic assignment revealed that bacterial and viral taxa dominated the microbial profiles, with occasional hits to fungal and other lineages (Fig. 5 A and B; Additional file 6). Bacterial composition included common *Drosophila* gut-associated taxa such as *Citrobacter*, *Fructilactobacillus*, and *Pseudomonas*. In addition, we found reads assigned to genera such as *Klebsiella*, *Escherichia*, *Streptococcus*, and *Salmonella*, which are rarely reported in *Drosophila* natural microbiome [Fig. 5 A and B; 48, 49]. Their presence here may reflect transient acquisition from rearing media, or lineage-specific associations unique to wild flies. Overall, these broad taxonomic patterns were generally similar between pathogen-challenged and unchallenged individuals within each species. However, more fine-scale differences emerged when we quantified within-sample microbial diversity (Fig. 5 C). Notably, *H. cameraria* exhibited a significant increase in alpha diversity following pathogen challenge (Wilcoxon rank-sum test, p = 0.01), suggesting that infection alters the richness or evenness of microbial communities in this species (Fig. 5 C). In contrast, microbial diversity remained stable across treatments in *H. confusa* and *S. deflexa* (Fig. 5 C). This increase in microbial diversity in *H. cameraria* may be associated with susceptibility to *Providencia* infection. Indeed, *Providencia* reads were more abundant in pathogen-challenged *H. cameraria* individuals, suggesting successful bacterial replication. In *H. confusa*, *Providencia* was detected at lower levels and in fewer individuals, and *S. deflexa* showed little evidence of *Providencia* infection, with only two challenged individuals containing detectable levels (Fig. 5 A). A few unchallenged individuals also showed low *Providencia* abundance, potentially due to environmental exposure or read contamination.

**Fig. 5:**
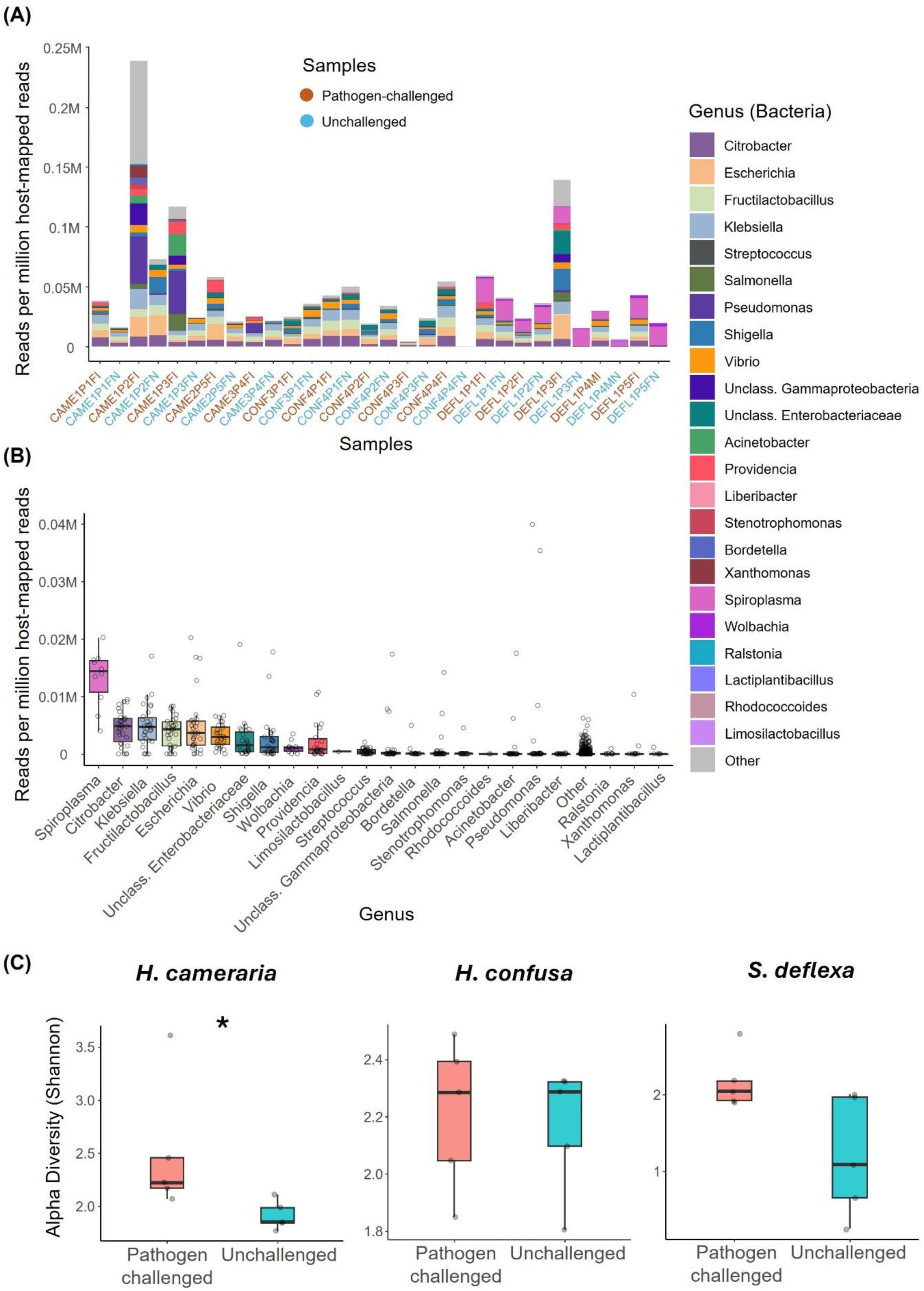
Microbiome composition and alpha diversity in pathogen-challenged and unchallenged samples across three drosophilid species. **(A)** Stacked bar plot showing the relative abundance of the top 23 bacterial genera (measured as reads per million host-mapped reads) across all samples. The 23 genera represent the union of the top 10 most abundant genera from each sample. Samples are ordered by species (*H. cameraria*, *H. confusa*, *S. deflexa*) and are color-coded by treatment status: pathogen-challenged (red) and unchallenged (blue). **(B)** Boxplot summarizing the relative abundance of each bacterial genus across all samples. Each box represents the distribution of abundance for a given genus, with individual data points indicating values from individual samples. Genera are ordered by median abundance. *Spiroplasma* shows the highest median abundance overall, and it’s only found in *S. deflexa*. **(C)** Boxplots of alpha diversity (Shannon index) comparing pathogen-challenged and unchallenged samples within each species. A significant increase in diversity is observed in *H. cameraria* upon infection (Wilcoxon rank-sum test, p value = 0.01), suggesting infection-induced shifts in microbial richness and/or evenness. No significant difference in alpha diversity was observed in *H. confusa* or *S. deflexa*.

Interestingly, we detected high levels of *Spiroplasma* in all *S. deflexa* individuals. This vertically transmitted bacterial endosymbiont is known to protect some species of *Drosophila* against parasitoids, nematodes, and bacterial pathogens [50–53]. This includes protection of *D. melanogaster* against *Providencia alcalifaciens* through host iron sequestration and enhanced melanization [53]. Crucially, these defense strategies can operate independently of canonical Toll and Imd pathway AMP gene upregulation while still providing effective physiological protection, and this may not result in (host) transcriptional signatures of a response to infection. *Wolbachia* was also detected at moderate levels in *S. deflexa*, though its effects are known to be more context dependent [54–56].

### Divergent species encode novel candidate AMP-like proteins

Despite the broadly conserved immune repertoire and similar transcriptional responses observed in *H. cameraria* and *H. confusa*, our analyses also revealed considerable species-specific differences—particularly in the apparently muted immune response of *S. deflexa*. These findings highlight the possibility that deeply diverged drosophilid species can encode lineage-specific immune effecters that are highly divergent in sequence and thus undetectable through simple homology searches against *D. melanogaster*. We hypothesized that such genes may include novel antimicrobial peptides, which are often short, secreted, cationic proteins with low levels of sequence conservation that makes them hard to detect [57, 58]. To identify potential novel AMP-like candidates, we focused on genes differentially expressed in response to pathogen challenge (adj. p < 0.05, log₂FC ≥ 1) that lacked detectable orthologs in *D. melanogaster*, and encoded short peptides of the length expected for known AMPs (i.e. ≤200 amino acids). Across the three species, this approach yielded a total of 41 candidates, 22 from *H. cameraria*, 13 from *H. confusa*, and 6 from *S. deflexa* (Additional file 7). Fourteen of these candidates were found in multispecies hierarchical orthogroups (HOGs), suggesting that a subset may represent conserved but previously unannotated drosophilid AMP families not found in *D. melanogaster*. The remaining candidates appeared to be species-specific or present in low-copy, potentially orphan gene families.

To evaluate their potential to encode AMPs, we screened all 41 candidates using two AMP prediction tools, AMP Scanner v2 [59] and amPEPpy [60], as well as SignalP 6.0 [61] for signal peptide prediction. Thirty-three of the 41 candidates were predicted as AMPs by at least one method, and 20 of these also had predicted N-terminal signal peptides (Additional file 7), indicating likely secretion. Based on physicochemical criteria, we further categorized the predicted AMPs into two groups: (i) Strong candidates (14) that encode positively charged (net charge +1 to +10), secreted peptides often enriched in glycine, proline, or cationic residues, and (ii) Likely candidates (6) that encode secreted peptides with weakly positive or negative net charge (–2 to +1), but are strongly induced by infection (Additional file 7). Among the strong candidates, we identified five in *H. cameraria*, seven in *H. confusa*, and two in *S. deflexa*. Expression levels of these genes were generally high (log₂ FC ranging from 1.08 to 8.30), and most were exclusively induced in pathogen-challenged individuals, supporting a role in infection response. Structural predictions using AlphaFold [62] revealed that many of these strong candidates adopt conformations typical of known AMPs, including amphipathic α-helices and αβ-sheet-rich peptides (Fig. 6).

**Fig. 6:**
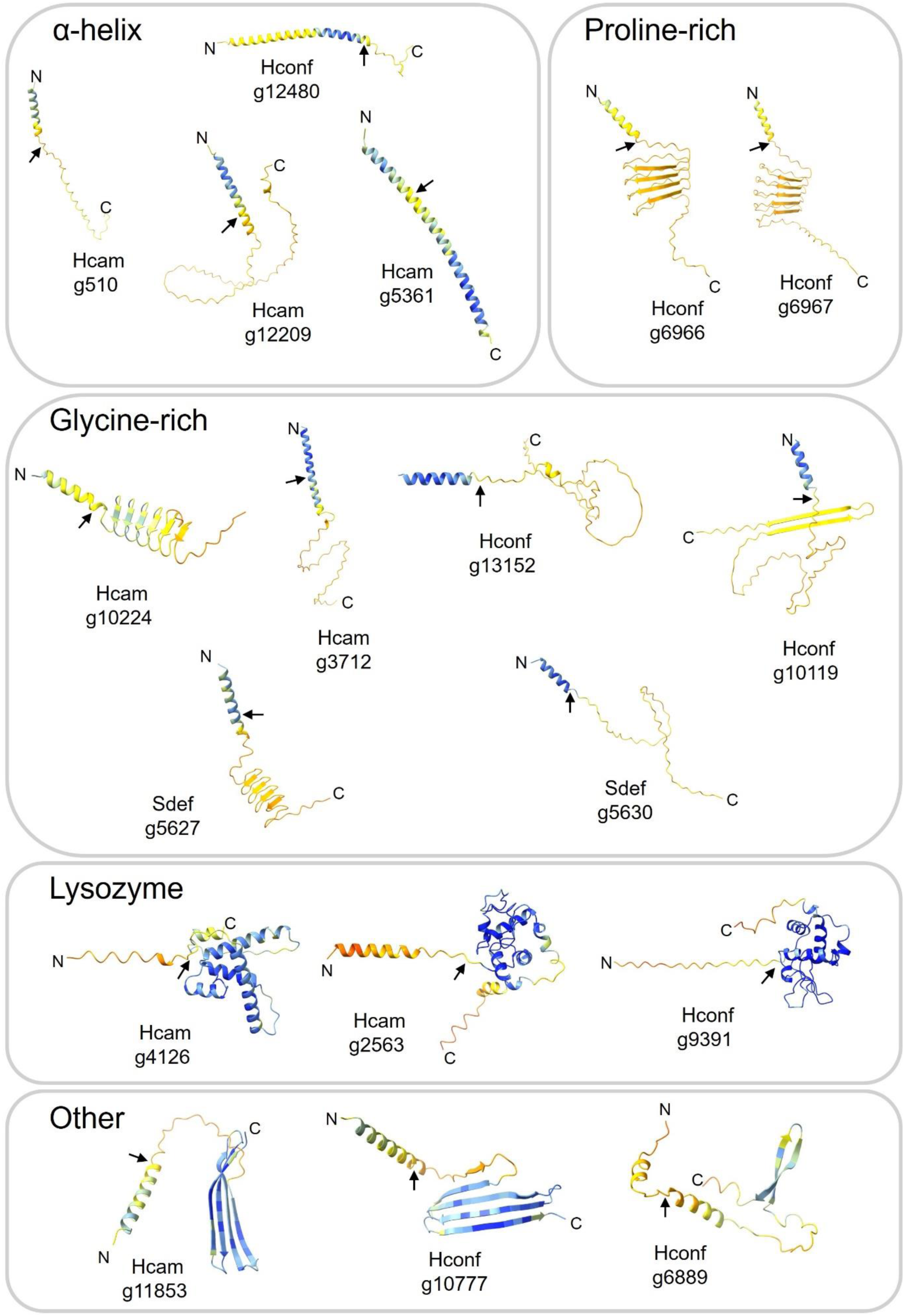
Predicted structures of novel candidate AMPs. AlphaFold3 predicted protein structures, N- and C-termini are labelled, and black arrows denote the predicted signal peptide cleavage sites based on SignalP6.0. The structures are colored by pLDDT confidence score (blue = high accuracy, light blue = backbone expected to model well, yellow/orange = lower confidence, red = very low confidence).

Several candidates exhibited striking similarity to well-characterized AMP classes. For example, the gene *Hconf/g6966* (∼20% proline) resembles the proline-rich *apidaecins* of bees [∼30% proline; 63], featuring four tandem repeats of potential mature peptides flanked by Furin cleavage motifs (RXRR). The gene *Hcam/g510* adopts a compact α-helical structure reminiscent of *IM18* [Paillotin; 64] from *D. melanogaster*, while *Sdef/g5630* (∼30% glycine) shows structural similarity to the glycine-rich AMP *holotricin* from *Aedes aegypti* [∼49% glycine; 65]. Additional candidates included *Hconf/g6967*, a paralog of *g6966*, also similar to *apidaecins*; *Hcam/g5361*, with ∼64% identity to *D. grimshawi CecC*; and *Hcam/g2563*, which shares ∼46% identity with *D. grimshawi lysozyme P*. Notably, both *Hcam/g4126* and *Hconf/g9391* have the potential to form *lysozyme*-like structures (Fig. 6). An especially intriguing example is *Hcam/g11853*, which was not predicted as an AMP by either amPEPpy [60] or AMP Scanner v2 [59], yet was among the most strongly induced genes following infection (log₂FC = 8.70; Table 2). Its homolog *Hconf/g10777* was similarly upregulated (log₂FC = 6.12) and clustered in the same HOG (Additional file 7). While blastp searches with this gene returned only uncharacterized *Drosophila* proteins, weak hits to the *edin* (‘elevated during infection’) gene family suggest that these genes may represent highly divergent *edin*-like effectors that escape homology detection—underscoring the evolutionary lability of this gene family in Drosophilidae. Predicted 3-D structures for all strong and likely AMP candidates are presented in Fig. 6. Taken together, their predicted secretion, AMP-like physicochemical properties, infection-induced expression, and structural similarity to known AMPs suggest that these genes likely encode novel immune effectors.

## Discussion

In this study, we provide a comparative analysis of transcriptomic responses following bacterial-challenge with Gram-negative pathogen *Providencia rettgeri* in three wild-derived, non-model drosophilid species—*H. cameraria*, *H. confusa*, and *S. deflexa*. These species, which diverged from *D. melanogaster* over 45 million years ago, represent largely uncharacterized and ecologically diverse branches of the drosophilid phylogeny. Our goals were to demonstrate the feasibility of differential expression analyses in near-wild flies, to assess whether canonical immune responses are conserved across these lineages, and to identify novel candidate immune effectors in taxa previously inaccessible to functional genomic studies. Through immune gene annotation, differential gene expression, microbiome profiling, and AMP prediction, our results reveal both conserved and lineage-specific features of insect immunity and highlight the value of including non-model species in comparative immune transcriptomic research.

Despite the intuitive appeal of using pathogen-stimulated transcriptomes to improve immune gene prediction in non-model species, we found that pathogen-challenged RNA-seq did not substantially improve annotation completeness or immune gene recovery. BRAKER3 based annotations derived from pathogen-challenged, unchallenged, and combined RNA-seq datasets yielded similar numbers of genes, with most orthologous genes consistently annotated across all datasets. While a few genes were unique to a single dataset, these rarely encoded immune-related functions and were often unassigned to orthogroups— suggesting lower confidence or assembly artefacts. These observations suggest that most immune genes have sufficient constitutive baseline expression to provide informative hints for de-novo gene prediction tools. Although RNA-seq also indirectly informs gene annotation by revealing new transcript isoforms and splicing events, and increases confidence, it is thus not essential for the identification of novel genes.

By identifying homologs of a curated set of 638 immune-related *D. melanogaster* genes, we found that all three species shared a conserved core of immune signaling and recognition genes. However, as in previous studies, we observed extensive lineage-specific variation among effector genes, particularly antimicrobial peptides, which were highly variable in presence, copy number, and inducibility [8, 18, 25]. Notably, *Scaptodrosophila deflexa* exhibited the lowest recovery of immune genes overall and lacked orthologs of several canonical AMPs, including *DptA*, *AttC*, and *AttD*. This pattern echoes previous finding in *Scaptodrosophila lebanonensis*, where a gene syntenic to *DptA* was identified but exhibited very little sequence similarity to any *diptericin* and in subgenus *Drosophila* is replaced by *DptC* [18]. It is therefore plausible that *DptA* has either been lost or has diverged beyond recognition in *S. deflexa*, potentially reflecting strong diversifying selection or relaxed constraint. In contrast, *H. cameraria* and *H. confusa* both harbored multiple paralogs of *DptC*, and *AttC*, suggesting lineage-specific expansions. These patterns of gain and loss were supported quantitatively by our Bayesian mixed-effects model, which showed that both species identity and gene functional category significantly predicted immune gene recovery. Effector genes were recovered with slightly, but significantly, lower probabilities than signaling genes, reinforcing the view that AMP gene families are particularly prone to evolutionary turnover. Among the species, *S. deflexa* showed the lowest predicted probability of immune gene recovery, again driven in part by the absence of some canonical AMP orthologs. These findings highlight the evolutionary lability of AMP gene families in Drosophilidae, characterized by frequent duplication, pseudogenization, and loss—hallmarks of strong, dynamic selective pressures likely imposed by pathogen diversity and host ecology. They also underscore the challenge of identifying fast-evolving immune effectors in non-model organisms, where high divergence can obscure orthology relationships and thereby our inability to functionally annotate these genes.

Interestingly, while both *H. cameraria* and *H. confusa* mounted robust transcriptional responses to *Providencia rettgeri* infection—upregulating canonical AMPs (such as *diptericins*, *attacins*, and *cecropins*), *PGRP* receptors, serine proteases, and transcriptional regulators (such as *Relish* and *pirk*)—*S. deflexa* appeared to exhibit minimal transcriptional changes, with only two *PGRPs* and no known AMPs upregulated. Assuming this was not simply lower power in our experiment, there are two likely explanations. First, *S. deflexa* individuals showed lower *Providencia* load, suggesting reduced infection burden or greater resistance. Second, all individuals harbored *Spiroplasma*, an endosymbiont known to protect flies from pathogenic bacteria (including *Providencia*) via mechanisms such as iron sequestration and melanization—defense strategies that bypass transcriptional AMP induction [53]. The concurrent presence of *Wolbachia* might further perturb the host immune responses, potentially by masking or dampening canonical transcriptional responses. In mosquitoes, experimental infection with *Wolbachia* protects against broad-spectrum anti-microbe and parasite by upregulating immune effector molecules and those involved in antimicrobial pathway [66–68]. Together, these findings suggest that both reduced *Providencia* infection success and the presence of defensive endosymbionts—especially *Spiroplasma*—may explain the weak immune transcriptional response observed in *S. deflexa*. More broadly, our results underscore the importance of profiling the microbiome when interpreting host transcriptional responses in non-model, wild-derived insects, where natural symbionts and background microbial variation may obscure or reshape immune phenotypes.

Finally, our identification of 41 novel AMP-like genes—many of which are not widely conserved across Drosophilidae, and are short, secreted, and structurally similar to known AMPs—suggests that AMP evolution may be even more dynamic than previously appreciated. Notably, many candidates shared structural features with known AMP families such as *apidaecins*, *holotricins*, *cecropins*, and *lysozymes*, but lacked clear sequence homology to any annotated *D. melanogaster* genes. Other candidates were not predicted to be AMPs, but were highly induced—including *Hcam/g11853* and its homolog *Hconf/g10777*, which could represent highly divergent homologs of *edin*. These results build on previous reports that lineage-specific AMPs are common in Drosophilidae [18, 25, 27] and emphasize that reliance on model species alone likely underestimates the diversity of immune effectors in nature.

### Conclusions

This study expands the scope of functional immune genomics into non-model drosophilids and highlights both the conserved and lineage-specific components of insect immunity, particularly we report extreme divergence and presence of novel AMPs in distant lineages of Drosophilidae. Our findings advocate for a broader phylogenetic sampling in immunological research and demonstrate the feasibility of high-resolution transcriptomics in species that are not amenable to laboratory domestication. In the future, the functional validation of novel immune effectors, through antimicrobial assays or in vivo perturbations will be critical to understand their roles in host defense. Additionally, as genome and transcriptome data continue to accumulate across Drosophilidae, studies like this will be essential for understanding how immune systems evolve, diversify, and interact with microbial environments across evolutionary time.

## Methods

### Fly collection and infection

We collected wild-mated female *Hirtodrosophila confusa* (n = 2), *H. cameraria* (n = 3 females), and *Scaptodrosophila deflexa* (n=1) from the Hermitage of Braid (Edinburgh, UK, all within 500m of 55.9 N, 3.2 W) on 4^th^ August 2023. Wild-caught female flies were housed separately under a 12:12-hour light-dark cycle and allowed to lay eggs. Wild caught *H. confusa* and *H. cameraria* were maintained on whole mushrooms (*Agaricus bisporus*), and *S. deflexa* on fermentative substrate intended to mimic sap flux conditions: *Drosophila* medium supplemented with tiny slices of ripe banana and cotton plug heads moistened with cider and maple syrup. Emerging first-generation female flies were collected and housed in same-sex vials containing standard Lewis medium [69] for 3–5 days prior to bacterial challenge. As a result, all flies were between 3 and 6 days old at the time of infection. For bacterial challenge, we used the ‘Dmel’ strain of *Providencia rettgeri* originally isolated from wild *Drosophila melanogaster* by Brian Lazzaro [70], and provided to us by Pedro Vale (University of Edinburgh). *Providencia rettgeri* cultures were initiated from a single colony and grown overnight in 10ml LB broth at 37°C with shaking. The bacterial culture was centrifuged at 5000 rpm for 5 min at 4°C and the supernatant was discarded. Bacteria were diluted to an optical density of OD₆₀₀ = 0.1 in sterile phosphate-buffered saline (PBS) prior to infection [71]. Flies were anesthetized on CO₂, and bacterial challenge was administered by puncturing the thorax with a 0.14 mm diameter stainless steel pin dipped in the bacterial suspension. Five females per species (four females and one male for *S. deflexa*) were challenged in this way. Control flies were anesthetized but otherwise left unmanipulated, and thus this experiment did not control for wounding effects. Both challenged and unchallenged flies were maintained at room temperature for 16 hours post-infection, then flash-frozen in Trizol reagent (Invitrogen) and stored at –80°C until RNA extraction. Extraction and sequencing were performed by Novogene sequencing facilities (www.novogene.com) using a TRizol extraction protocol and Illumina NovaSeq sequencing platform to generate 150bp paired-end reads.

### Quality control and mapping

On average, samples contained 58.35 million raw reads per fly. We preprocessed raw reads for quality control using fastp v0.24.0, with the “*-c*” option enabled for overlap base correction and “*-y -Y 20*” for low-complexity filtering [72]. All other parameters were kept at their default settings. Reads were mapped to their respective genomes using STAR RNA-seq aligner v2.7.10b, generating sorted BAM files as output [73]. On average, 83.98%, 66.55%, and 82.02% of reads per library mapped uniquely to the *H. confusa* (assembly accession: GCA_035043065.1), *H. cameraria* (GCA_949708635.1), and *S. deflexa* genomes, respectively (Additional file 8). The *Scaptodrosophila deflexa* genome was sequenced by Bernard Kim (Princeton) and Dmitri Petrov (Stanford), using ONT R10.4.1 sequencing technology and assembled with hifiasm [74], and was generously made available to us in advance of publication. BUSCO completeness and N50 for all the genomes used in this study can be found in Additional file 8.

### Gene annotation

To assess the impact of pathogen challenged RNA-seq on our ability to recover immune genes, as compared with unchallenged RNA-seq, we generated 3 independent genome annotations for each species using: (1) RNA-seq data from pathogen-challenged individuals, (2) RNA-seq data from unchallenged (naïve) individuals, and (3) combined RNA-seq data. Genome annotations were generated using BRAKER3 in ETP mode, with extrinsic protein hints provided from *D. melanogaster* RefSeq proteins [38]. To assess gene orthology and recover immune-related orthogroups, we extracted the longest isoform per gene from each annotation set and ran OrthoFinder v2.5.5 using the predicted proteomes of all species and annotation sets, along with the *D. melanogaster* proteome [75]. To generate hierarchical orthogroups (HOGs), gene trees were inferred by OrthoFinder using alignments generated with MAFFT [76] and maximum-likelihood inference performed with IQ-TREE2 [77].

### Differential gene expression analysis and functional annotation

Read counts per gene were quantified using “featureCounts” [78]. Genes with fewer than 10 total counts across all samples were excluded from downstream analysis. Differential expression analysis was conducted using the DESeq2 package in R [42]. We performed principal component analysis (PCA) using the *plotPCA()* function on regularized log-transformed (rlog) counts, with *∼treatment* specified as the design formula. The top 500 most variable genes were used to calculate principal components. Genes were considered significantly differentially expressed if they had an adjusted p-value < 0.05 and a |log₂ fold change| ≥ 1. Heatmaps of expression patterns were generated using the pheatmap package.

Functional annotation of genes was performed using eggNOG-mapper v2.1.12 with Diptera HMM database using HMMER searches (*-m hmmer*) and additional filters for hits with e-value ≤ 0.05, bit score ≥ 60, and percent identity ≥ 40 [79, 80]. GO terms were restricted to non-electronic evidence codes, and PFAM domains were realigned (--pfam_realign realign). Gene Ontology (GO) enrichment analysis was performed using “topGO” package in R by applying Fisher’s exact test and the “weight01” algorithm, which accounts for the hierarchical structure of GO terms [81]. The p-values from GO analyses were corrected using the Benjamini and Hochberg procedure with the FDR threshold set to 0.05 [82].

### Metagenomic analysis of unmapped reads

To quantify microbial abundance in flies and to assess whether pathogen-induced expression differences could be affected by microbial load, we performed de novo metatranscriptomic analysis on unmapped reads from each sample. Paired-end unmapped reads were extracted from STAR-mapped BAM files using “samtools” v1.13 with the flags *-f 12 -F 256*, to retain only unmapped read pairs [83]. These reads were assembled de novo using “rnaSPAdes” v4.1.0 [84], with default parameters. Open reading frames (ORFs) were predicted from each transcriptome assembly using EMBOSS “getorf”, retaining protein length ≥200 amino acids. The resulting proteins were queried against the NCBI non-redundant (nr) protein database using “diamond blastp*”* v2.1.10 (*--evalue 1e-20, --outfmt 6*) [47] and taxonomic lineages were assigned to diamond hits using “TaxonKit” [85]. To estimate relative microbial load, total reads mapped to each organism were normalized to host-mapped read counts for each sample to control for sequencing depth. Alpha diversity (Shannon index) was calculated from genus-level profiles using the vegan and phyloseq packages in R [86, 87]. Statistical comparisons between pathogen-challenged and unchallenged groups were performed using the Wilcoxon rank-sum test.

### Prediction of novel AMPs

To identify potential novel immune effectors, we screened all significantly upregulated genes that had no detectable homologs in *D. melanogaster* and a predicted peptide length ≤ 200 amino acids. These candidates were assessed using two AMP prediction tools: (1) AMP Scanner v2, a deep neural network-based classifier [59], and (2) amPEPpy, a Python implementation of the amPEP random forest classifier [60]. The random forest model was trained using 712 known AMPs from the APD3 database and 712 matched non-AMP sequences [see original publication for dataset details: 59]. All candidate peptides were also analyzed for the presence of N-terminal signal peptides using SignalP 6.0 webserver [61]. Physiochemical properties of candidates were obtained using AMP predictor tool of APD3 [88] and Expasy “ProtParam” [89]. Finally, 3-D structures of AMP candidates were generated using the AlphaFold 3 server [62].

## Availability of data and materials

All RNA-seq reads data generated for this study have been deposited in the SRA database, and can be found under project ID PRJNA1270041.

## Acknowledgements

We would like to thank the Friends of the Hermitage of Braid and City of Edinburgh Forestry and Natural Heritage for permission to collect flies. We also would like to thank Bernard Kim and Dmitri Petrov for the pre-publication *Scaptodrosophila* genome, and Katy Monteith and Pedro Vale for providing the bacterial aliquots. For the purpose of open access, the authors have applied a Creative Commons Attribution (CC BY) license to any Author Accepted Manuscript version arising from this submission.

## Funding

This work was supported by a Darwin trust PhD studentship to PD and Biotechnology and Biological Sciences Research Council grant BB/T007516/1 to DJO.

## Notes

### Competing Interest Statement

The authors have declared no competing interest.

