## Supplementary figures and images for "Transcriptomic analysis of non-model Drosophilidae reveals novel AMP candidates"

### Additional file 1

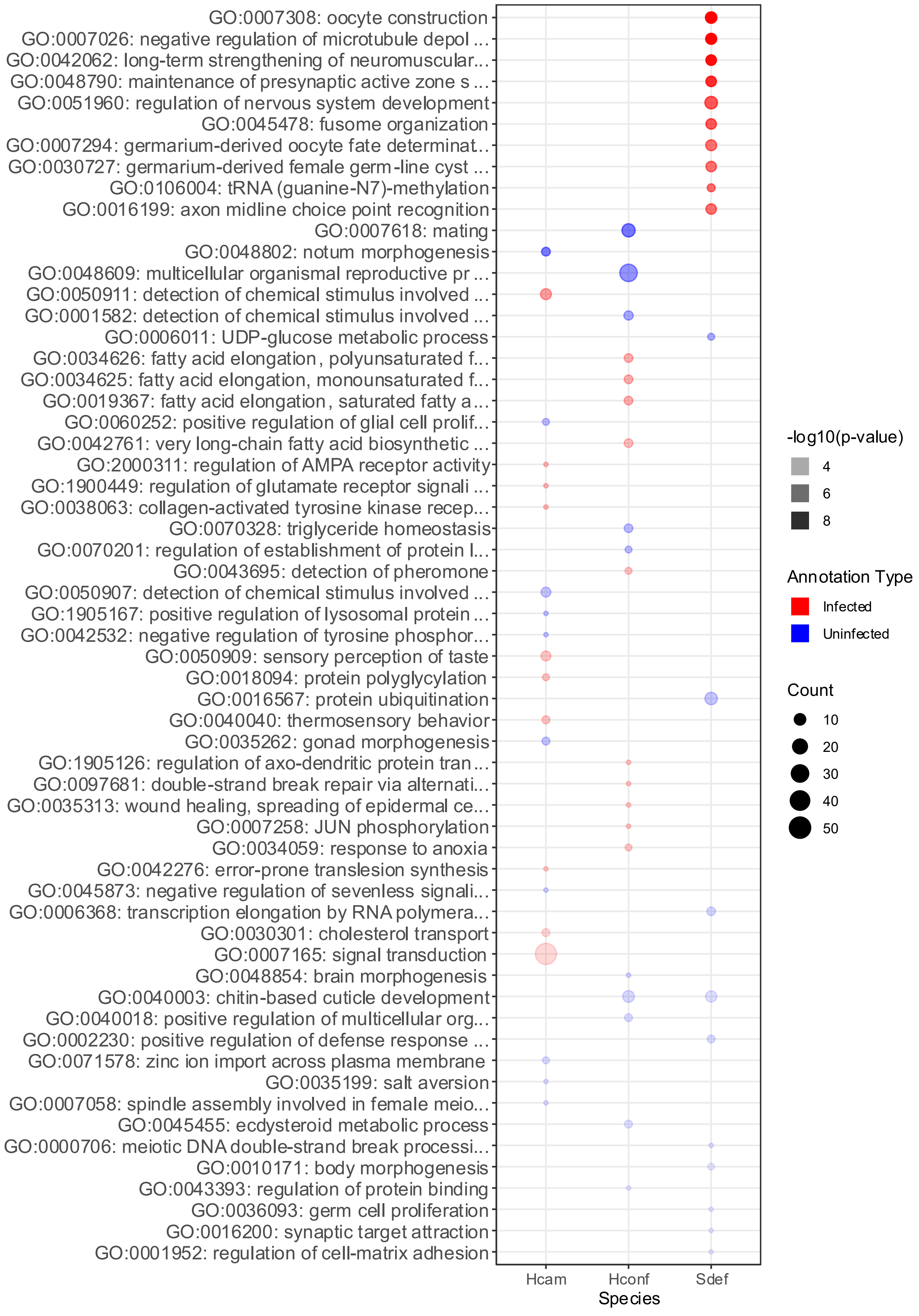

### Additional file 3

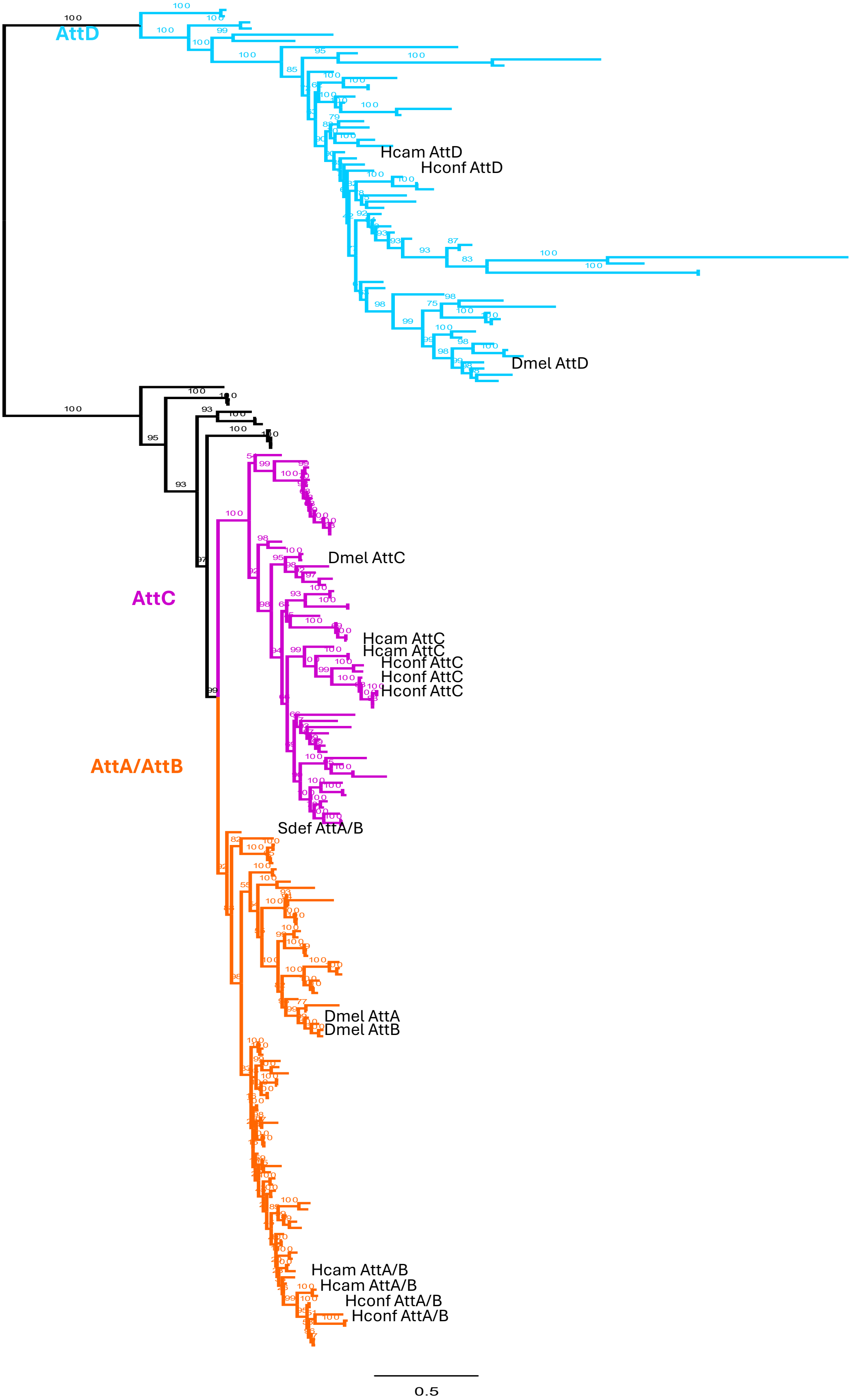

### Additional file 5

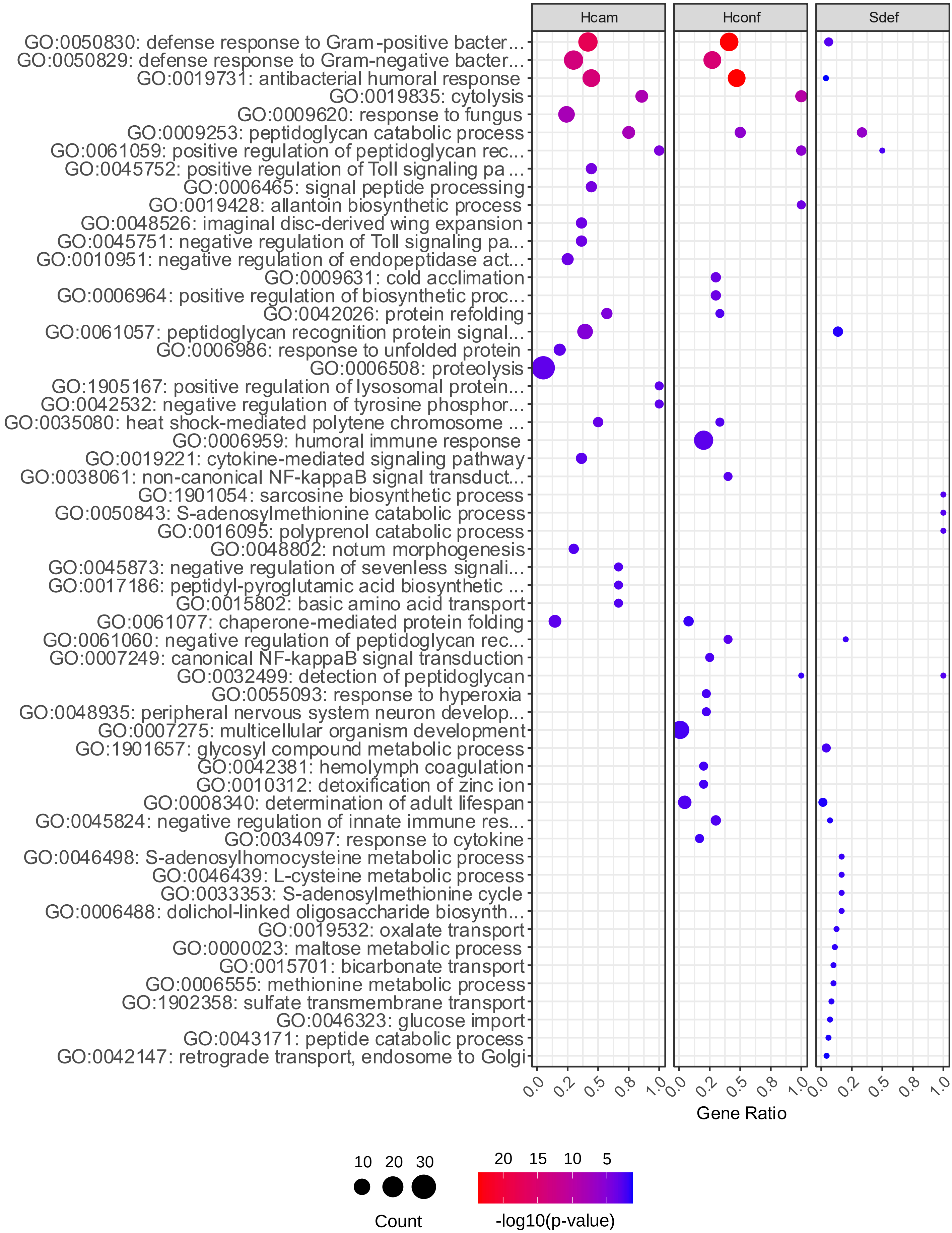

### Additional file 6

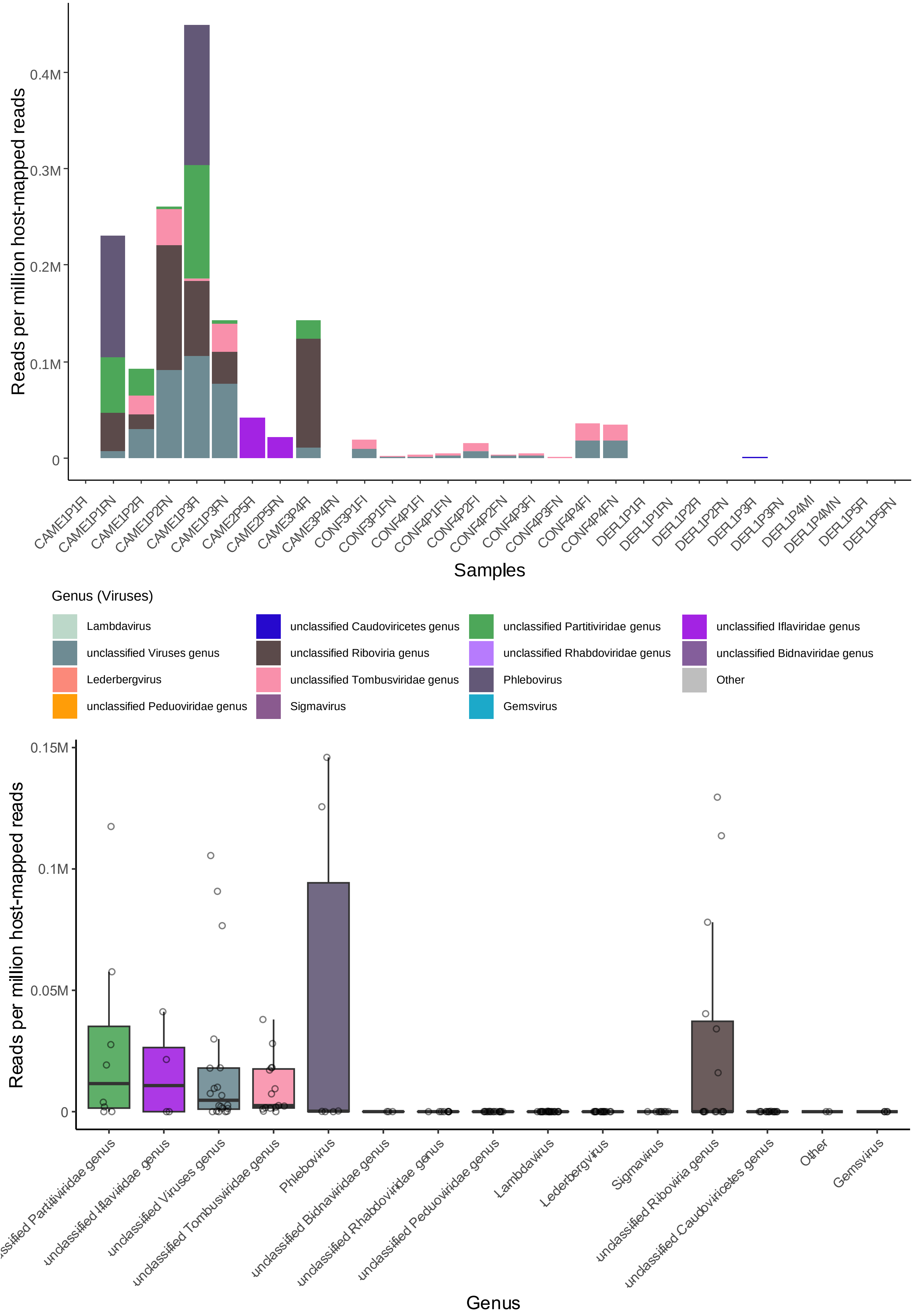
